# Downregulation of hippocampal *NR2A/2B* subunits related to cognitive impairment in a pristane-induced lupus *BALB/c* mice

**DOI:** 10.1101/631879

**Authors:** Jonatan Luciano-Jaramillo, Flavio Sandoval-García, Mónica Vázquez-Del Mercado, Yanet Karina Gutiérrez-Mercado, Rosa Elena Navarro-Hernández, Erika Aurora Martínez-García, Oscar Pizano-Martínez, Fernanda Isadora Corona-Meraz, Jacinto Bañuelos-Pineda, Jorge Floresvillar-Mosqueda, Beatriz Teresita Martín-Márquez

## Abstract

Neuropsychiatric systemic lupus erythematosus (NPSLE) is a severe complication associated with the neurotoxic effects of circulating autoantibodies in the central nervous system (CNS) manifested frequently as a learning and memory deficit. Pristane-induced lupus in *BALB/c* female mice is an experimental model that resembles some clinical and immunological SLE pathogenesis associated with environmental factors. Nevertheless, there is no experimental evidence that relate pristane-induced lupus with cognitive dysfunction associated with autoantibodies production.

**Objective:** To evaluate cognitive impairment related to memory deficits in a pristane-induced lupus *BALB/c* female mice related to mRNA expression levels of *NR2A/2B* hippocampal subunits in short and long-term memory task at 7 and 12 weeks after LPS exposition (7wLPS and 12wLPS) in a behavioral test with the employment of Barnes maze.

**Methods:** Fifty-four female *BALB/c* mice of 8-12 weeks old were included in 2 experimental groups: 7 and 12 weeks after lypopolissacharide (LPS) exposure and classified in subgroups (control, pristane and pristane+LPS). To determine cognitive dysfunction, mice were tested in a Barnes maze. Serum anti-Sm antibodies and relative expression of hippocampal *NR2A/NR2B* subunits were quantified.

**Results:** Pristane and pristane+LPS mice showed a prolonged escape latency at 7wLPS than at 12wLPS in short-term memory. Downregulation of hippocampal *NR2A* subunit was more evident than *NR2B* in pristane and pristane+LPS at 7wLPS and 12wLPS. The anti-Sm autoantibodies levels correlate with the relative expression of *NR2A.*

**Conclusion:** Downregulation of hippocampal *NR2A/2B* subunits in the pristane-model of lupus in *BALB/c* mice may be related to anti-Sm autoantibodies production with the consequence of cognitive impairment in early stages of autoimmune disease.

## Introduction

Systemic lupus erythematosus (SLE) is an idiopathic autoimmune disorder characterized by induction of autoantibodies against intracellular components such as nucleosomes (double-stranded DNA and histones) and small nuclear ribonucleoproteins (snRNPs) known as Smith antigen (anti-Sm), that is consider for the American College of Rheumatology (ACR) as a classification criteria for SLE diagnosis [1]. This condition presents a wide variety of clinical manifestations with multiple organs affectations, especially skin and kidneys, however, the heart and nervous system are also damaged [2]. In SLE, central and peripheral nervous system are involved in the development of psychiatric abnormalities termed neuropsychiatric-SLE (NPSLE) syndromes [3]. In 1999, the ACR established a standard nomenclature with case definitions for 19 neuropsychiatric conditions, 12 related to central nervous system (CNS) manifestations (mainly seizures, headache, stroke, depression, cognitive dysfunction, and psychosis) [3, 4]. Clinical studies estimate an NPSLE prevalence in a range from 17 to 80%, these variations can be attributed to diagnostic criteria, patient selection and assessment methods for autoantibodies detections [3, 5–7]. The etiopathogenesis of NPSLE is still unknown, however, several studies suggest that the presence of autoantibodies against to N-methyl-D-aspartate (NMDA) receptors in serum and cerebrospinal fluid (CF), the production of intrathecal proinflammatory cytokines/chemokines and vasculitis are associated to neuropsychiatric manifestations such as cognitive dysfunction [2, 3]. Analysis in SLE patients and murine model of lupus report that certain subsets of anti-double-stranded DNA (anti-dsDNA) and anti-NMDA transit from the vasculature to the amygdala when the blood-brain barrier (BBB) permeability was altered and cross-react with a consensus pentapeptide (DWEYS) present in NR2A and NR2B subunits of NMDA receptor, mediate neuronal loss and affects learning and memory [2, 8–10].

To reproduce clinical and molecular NPSLE physiopathology, there have been developed experimental models of lupus brain diseases, and to be consider a brain model and resemble the neuropathology condition, they must gather two requirements: the production of autoantibodies that cross-react with neuronal receptors and the disruption of BBB integrity by exposure to lipopolysaccharide (LPS) [2]. Lupus can be induced by exposure a healthy mouse strain (*BALB/c*) to hydrocarbon oils such as pristane (2,6,10,14-tetramethylpentadecane) that induce a wide range of specific SLE autoantibodies (anti-DNA, anti-RNP/Sm and anti-Su) in a range between 12 to 25 weeks in the trial period [11–13]. This is a suitable model to evaluate the break tolerance related to environmental factors associated with SLE development. Nevertheless, there are no experimental evidences that relate pristane-induce lupus with cognitive dysfunction associated with the development of autoantibodies against hippocampal NMDA receptor subunits NR2A/2B.

In order to evaluate cognitive impairment related to memory deficits in a pristane-induced lupus *BALB/c* mice, we analyzed the mRNA expression levels of *NR2A/2B* hippocampal subunits in short and long-term memory task at 7 and 12 weeks after LPS exposition with a behavioral test with the employment of Barnes maze.

## Materials and methods

### Animals

Female *BALB/c* mice of 8-12 weeks old were obtained from UNAM-Envigo RMS Laboratory in México City and housed in the animal facility of Instituto de Investigación en Reumatología y del Sistema Músculo Esquelético of Centro Universitario de Ciencias de la Salud under the following conditions: 2-4 animals in clear cages (7.6×11.6×4.8 inches), controlled temperature room at 22±1°C, positive laminar flow, 12 hours of light/dark cycles and feed *ad libitum* with purified water and normocaloric diet (Rodent Chow 5001, Purina™). The protocol was approved by the Committee of Investigation, Ethics and Biosecurity of Centro Universitario de Ciencias de la Salud of the University of Guadalajara (Protocol number CI-07918) and all experimental procedures were carried out in compliance with the Rules for Research in Health Matters (Official Mexican Norms NOM 0062-ZOO-1999 and NOM-033-ZOO-1995).

### Induction of lupus by pristane and LPS exposure

A total of 54 female *BALB/c* mice of 8-12 weeks old were included and separated into 2 experimental groups: 7 and 12 weeks after LPS exposure (abbreviated as 7wLPS and 12wLPS respectively), and in a subgroups denominated control (single intraperitoneal injection (i.p.) of 0.5 mL NaCl 0.9%), pristane (single i.p. pristane injection, Sigma Chemical Co, St Louis, MO, USA) and pristane+LPS (single i.p. pristane injection and LPS of *E. coli* O55:B5, Sigma St Louis, MO, USA in a dose of 3mg/kg diluted in NaCl 0.9% 16 weeks post-pristane administration [14]). The 7wLPS group were integrated by 8 controls, 10 pristane, and 10 pristane+LPS and for 12wLPS group we included 6 controls, 10 pristane and 10 pristane+LPS (Fig 1, A).

**Fig 1.**
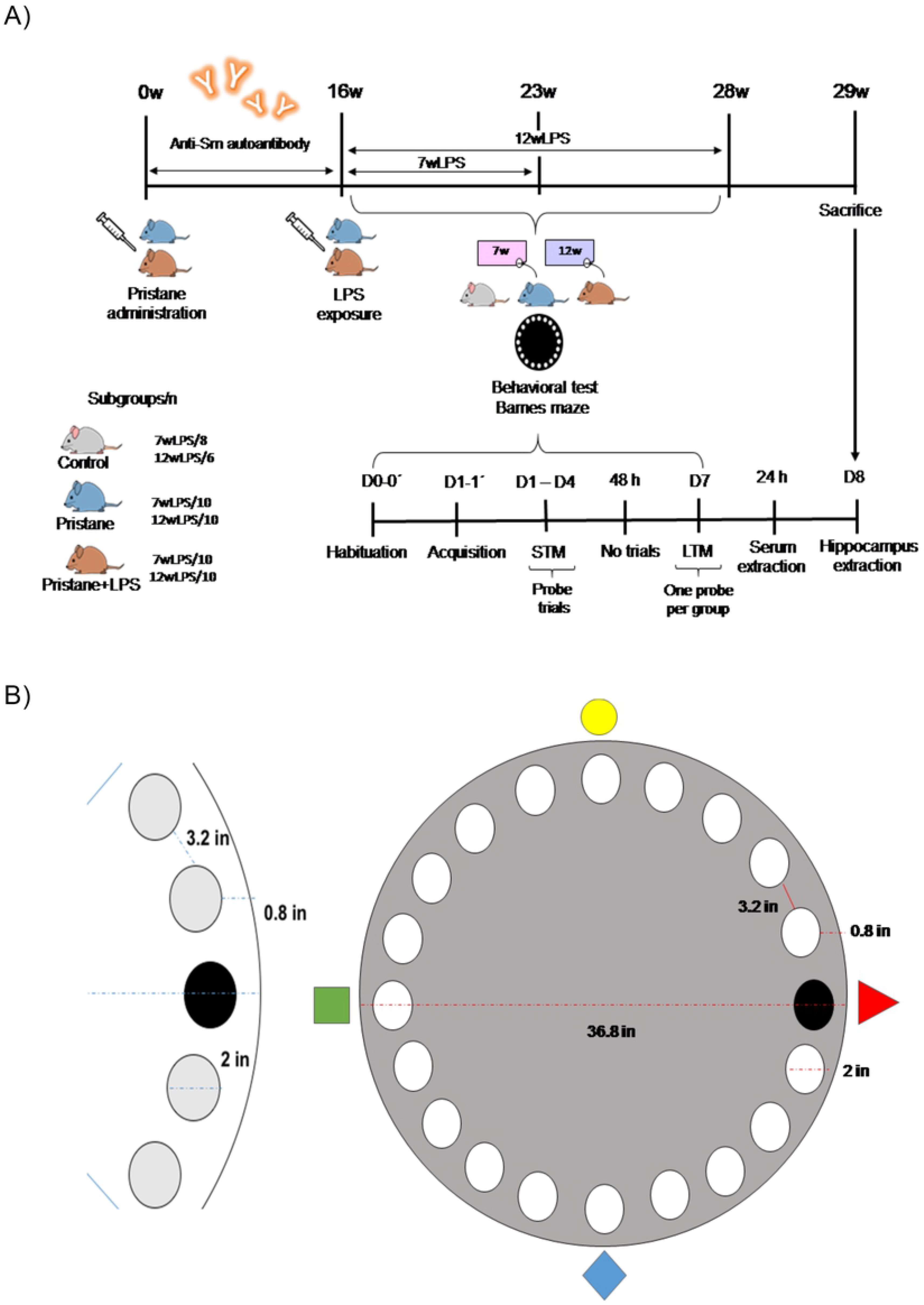
Experimental procedures squeme and Barnes maze. A) Time schedule of experimental procedures. B) Barnes maze platform designed for mouse behavioral testing.

### Barnes maze

To assess memory and learning process to determine cognitive dysfunction, experimental groups of 7wLPS and 12wLPS were evaluated in a behavioral test with a Barnes maze adapted for mouse based on the protocol described by Barnes *et al* [15]. For the maze, we employed a circular black acrylic platform of 36.8 inches of diameter anchored in a metallic base of 32 inches of height above the floor. The platform has 20 holes of 2 inches of diameter disposed along the perimeter with 3.2 inches among them (19 empty holes and one scape hole with dark box). For reference points to reach the escape hole, we used extra-maze cues around the room (circle, rhombus, triangle, and square in different colors) and for eliminating odor cues, the experimentator cleaned the platform and the scape box in every trial with ethyl alcohol at 70% (Fig 1, B). To record the cognitive performance, we used a video tracking system (Pro Webcam-C920 HD 1080p).

### Behavioral test

The behavioral tests were developed by an experimentator in three phases: habituation, acquisition and probe trial assessed in an airy and odor free white room without visual and sound distractors. The habituation consisted in two days (Day 0 and Day 0’) with two trials per mouse into the platform and escape hole for a lapse of 180 seconds. For the acquisition phase (Day 1 and Day 1’) we included two trials per mouse and consisted in place the mouse into a white acrylic cylindrical chamber in the middle of the platform for 10 seconds and then released it to explore the platform for a lapse of 180 seconds, finishing the trial when the mouse entered by itself into the escape hole. In this phase, if the mouse did not enter in the target hole, it can be guided by the experimentator. The probe trials were assessed to evaluate memory and learning consolidation in mice that conformed subgroups and consisted in three repetitions per mouse on the maze to reach the escape hole for 180 seconds, taking in account all the procedures described previously. This phase was denominated Short Time Memory (STM) and consisted of four probe trials (D1-D4). Once finished, mice were preserved for a lapse of 48 hours and then evaluated for the Long Time Memory (LTM) at Day 7. We considered to evaluate the behavioral performance with two parameters: the latency in seconds to reach and enter in the escape box and the errors established as the deflections of the head in empty holes for each mouse prior to find and enter in escape box (Fig 1A).

### Anti-Sm antibody ELISA

Once the behavioral tests were finished, we obtained total blood from the tail vein of each mouse, the serum was separated and stored at −20°C. Levels of mouse anti-Sm antibodies in sera from experimental groups were measured by enzyme-linked immunosorbent assay (ELISA) using the quantitative kit Mouse Anti-Sm Ig’s (total/A+G+M) (Alpha Diagnostic International™) in a 1:2 dilution based on reference standards provided by the manufacturer.

### RNA isolation and *NR2A/2B* subunits qRT-PCR analysis

Once finished the behavioral test, mice were euthanized by CO_2_ inhalation and after craniotomy surgery, the hippocampus region was removed to obtain lysates for total RNA isolation. This procedure was performed according to the manufacturer’s procedure using the GF-1 Total RNA extraction kit (Vivantis Technologies™). The complementary DNA synthesis (cDNA) was performed with 5μg of each total RNA sample using a reaction size of 20μL with oligo (dT) primer (100 ng/μL), RNase free, DEPC treated water and Moloney Murine Leukemia Virus Reverse Transcriptase (M-MLV RT) kit (Applied Biosystems, 850 Lincoln Centre Drive, Foster City, CA 94404) and store at −20°C until being used for expression analysis. Real-time quantitative polymerase chain reaction (qRT-PCR) was conducted using Rotor-Gen (Q5 PLEX HRM System, QiagenTM) and the Q-Rex software was used for the analysis. A threshold cycle (C_T_) value was determined from each amplification plot. For *Mus musculus* genes, the specific primers were synthesized based in sequences published by Hamada *et al.*[16] as follows: *GluN2A* forward 5’-CCTTTGTGGAGACAGGAATCA-3’ and reverse 5’-AGAGGCGCTGAAGGGTTC-3’; *GluN2B* forward 5’-GGGTTACAACCGGTGCCTA-3’ and reverse 5’-CTTTGCCGATGGTGAAAGAT-3’. Expression of target genes was normalized with the endogenous reference mouse gene *GAPDH* using the following primers: forward 5’-TGTCCGTCGTGGATCTGAC-3’ and reverse 5’-CCTGCTTCACCACCTTCTTG-3’. The qPCR was performed in a final reaction volume of 10μL (10μM forward and reverse primer, 25μM ROX, 2x SYBR Green qPCR master mix and 100ng cDNA). The conditions of reaction were: holding at 95°C/10 min, cycling at 35 cycles of 95°C/10s and 55°C/15s and melt curve at 95°C/15s, 72°C/60s, and 95°C/15s.

### Statistical analysis

Comparisons were made using Kruskal-Wallis, post hoc tests were carried out using Mann-Whitney U as applicable. Values are presented as median with percentile 25 and 75 (P_25_-P_75_) and as mean and standard error of median (±SEM), as applicable. Spearman’s correlations coefficients were also calculated. All data were analyzed using SPSS v22.0 (SPSS Inc. Chicago, IL) and GraphPad Prism version 6.00 for Windows (GraphPad Software, La Jolla, CA). *P* <0.05 was considered statistically significant.

## Results

### Pristane and pristane+LPS treated mice showed a prolonged escape latency in STM at 7wLPS

Once the mice completed the habituation and acquisition probes in Barnes maze, we evaluated the STM in control, pristane and pristane+LPS subgroups at 7wLPS during D1-D4 (Fig 2). In D1, we observed differences in the exploration time to reach the target hole and enter in escape box between pristane 162.2s (120-180s) *vs.* pristane+LPS 180s (179.9-180s, *P*=0.033); in D2 control 97.3s (34.7-133.6s) *vs.* pristane+LPS (113-180s, *P=* 0.009); in D3 in control 22.1s (14.9-30.1s) *vs.* pristane 125s (33.3-180s, *P*= 0.014) and control *vs*. pristane+LPS 145.6s (116.5-152.8s, *P*<0.0001) and in D4 between control 12.2s (7.0-39.1s) *vs*. pristane+LPS 175s (40.3-180s, *P*=0.003). We were able in this first test to distinguish a different behavioral pattern between control and subgroups of pristane-treated mice. In this point, it is important to highlight that the subgroup of pristane and pristane+LPS showed in D1 of the probe the same behavioral pattern than the control subgroup and as the test was progressed, the mice treated with pristane and pristane+LPS showed an erratic behavior observable in D3-D4 and attributable to deficient memory retention.

**Fig 2.**
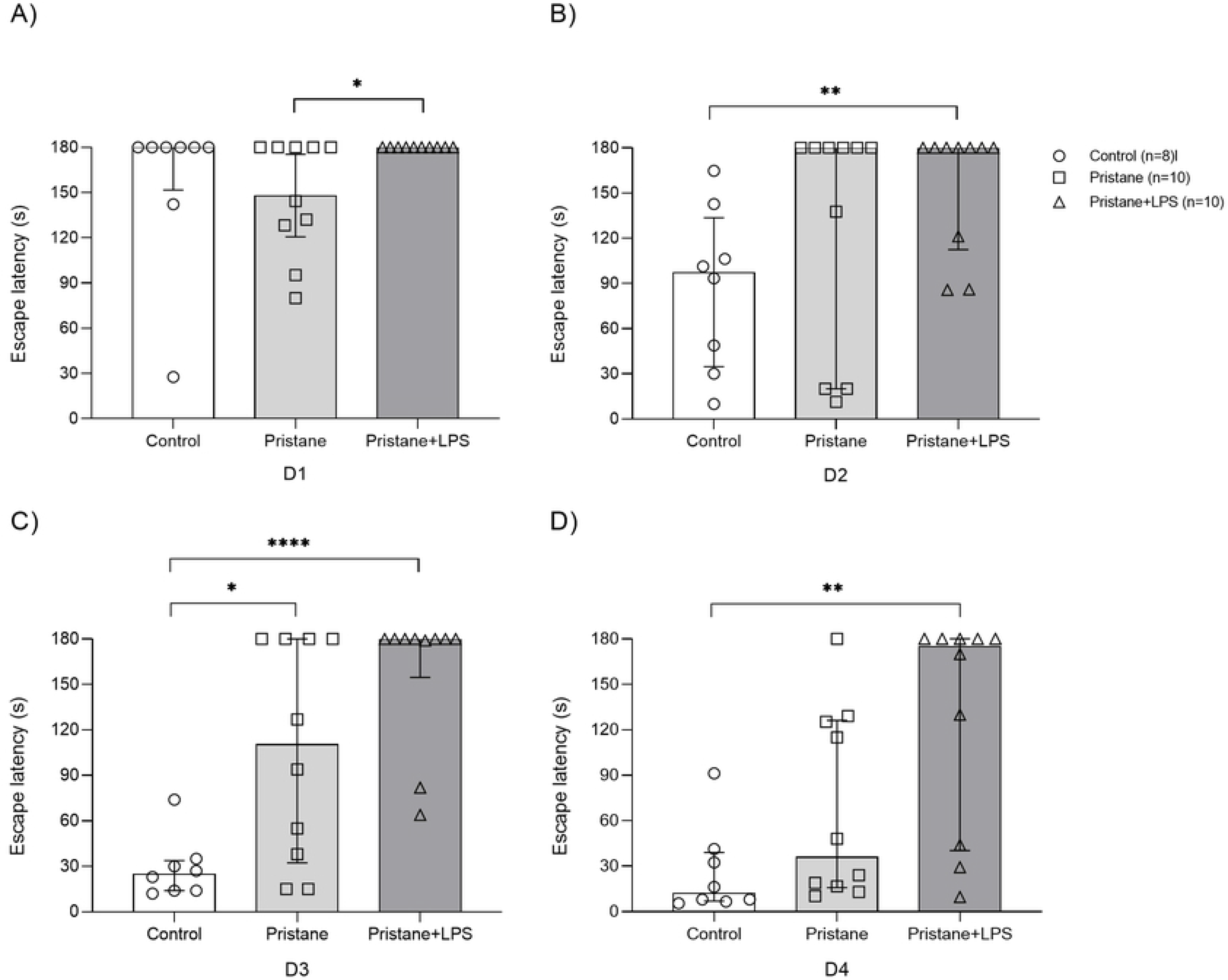
Escape latency of experimental subgroups in STM at 7wLPS. Total time average in seconds (s) of subgroups control, pristane and pristane+LPS to reach and enter to scape box during D1-D4 of STM. A) D1, B) D2, C) D3, D) D4. Values are presented as median with percentile 25 and 75 (P_25_-P_75_). * *P*<0.05, ** *P*<0.01, \*\*\*\**P*< 0.0001. At the same time of STM evaluation, we calculated the number of errors between groups in D1-D4 (S1 Fig) and we observed differences only in D4 between control 0.5 (0-1) *vs*. pristane 3 (2-6, *P*=0.04).

**S1 Fig. Errors in STM at 7wLPS.** Total errors of subgroups in STM at 7wLPS. A) D1, B) D2, C) D3, D) D4. Values are presented as median with percentile 25 and 75 (P_25_-P_75_). ** *P*<0.01.

### Pristane and pristane+LPS treated mice showed a prolonged escape latency in STM at 12wLPS

With the purpose to determine if the induction of lupus by pristane and the exposure LPS maintain its effects in cognition in a prolonged manner, we used the same experimental subgroups and evaluate them at 12wLPS (Fig 3). We observed statistical difference in escape latency only in D2 between control *vs.* pristane (141.5s *vs.* 180s, *P*=0.013) and control *vs.* pristane+LPS (141.5s *vs*. 180s 171.5-180, *P*=0.013) and although there were no differences in D1 and D3-D4, we observed that some mice of pristane and pristane+LPS subgroups showed prolonged escape latency in D3-D4.

**Fig 3.**
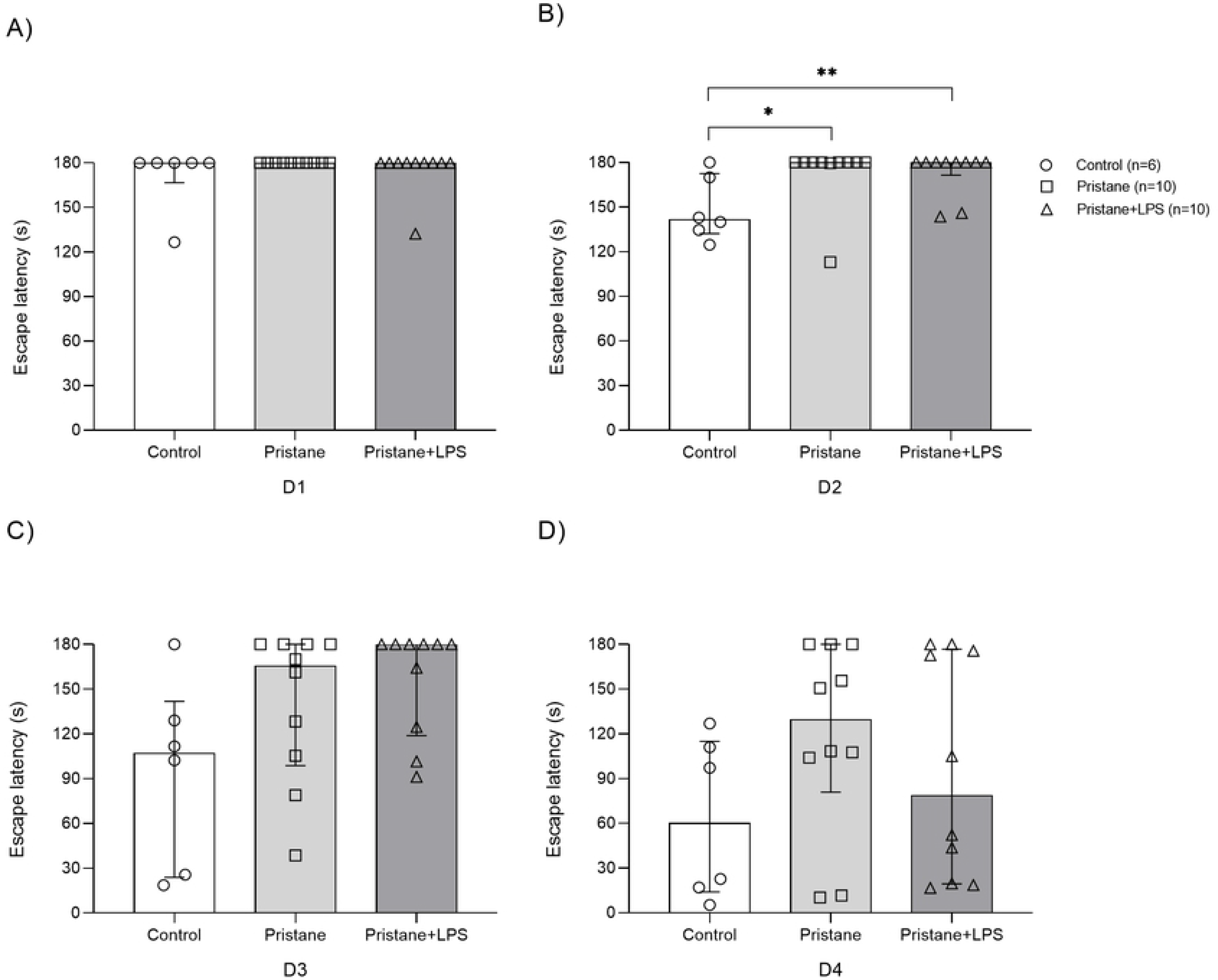
Escape latency of experimental subgroups in STM at 12wLPS. Total time average in seconds (s) for each mouse to reach and enter to scape box during D1-D4 of STM at 12wLPS. A) D1, B) D2, C) D3, D) D4. Values are presented as median with percentile 25 and 75 (P_25_-P_75_). \**P* < 0.05, \*\**P*<0.01.

During the test, the errors were calculated and we did not observe differences between the subgroups during D1-D4 in STM at 12wLPS (S2 Fig).

**S2 Fig. STM errors at 12wLPS.** Total errors of subgroups at 12wLPS during D1-D4. A) D1, B) D2, C) D3, D) D4. Values are presented as median with percentile 25 and 75 (P_25_-P_75_).

### Pristane and pristane+LPS treated mice showed a prolonged scape latency in LTM at 7wLPS

To determine learning and memory consolidation, we evaluated LTM 48 hours after STM test in 7wLPS subgroups and we observed a prolonged escape latency between control *vs.* pristane (6s *vs.* 14s, *P*=0.005) and control *vs*. pristane+LPS (6s *vs.* 35s, *P*<0.0001). In this probe, the total of control mice reaches the target hole in less than seconds in comparison to mice of pristane and pristane+LPS subgroups (Fig 4).

**Fig 4.**
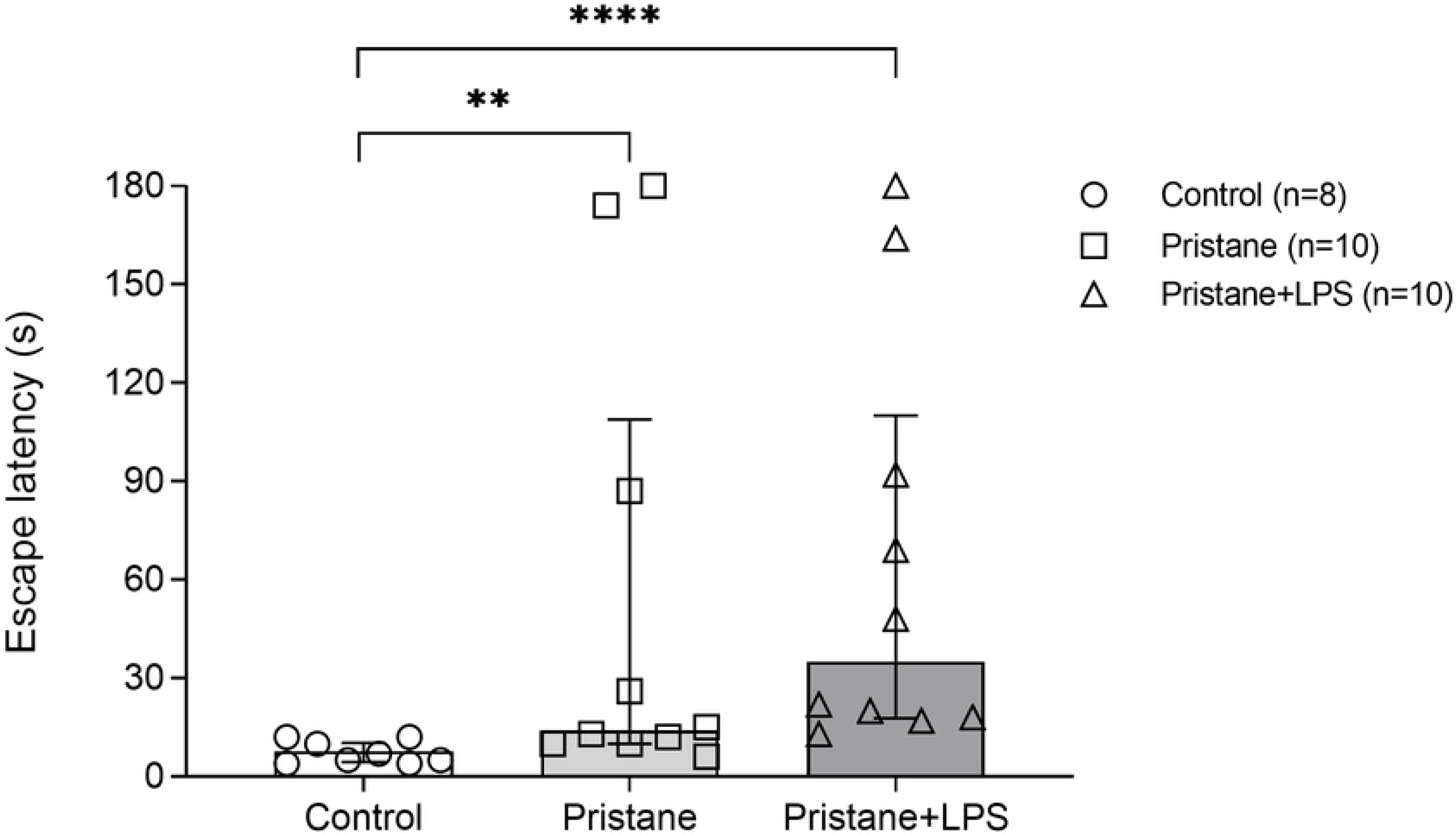
Escape latency of experimental subgroups in LTM at 7wLPS. Total time average in seconds (s) for each mouse to reach and enter to escape box during LTM at 7wLPS. A) D1, B) D2, C) D3, D) D4. The data were shown in medians and percentile 25 and 75 (P_25_-P_75_). \*\**P*<0.01, **** *P*<0.0001.

In relation to the errors in this probe, we did not observe differences between subgroups (S3 Fig).

**S3 Fig. LTM errors at 7wLPS.** Total errors of subgroups at 7wLPS in LTM probe. Values are presented as median with percentile 25 and 75 (P_25_-P_75_).

### Pristane and pristane+LPS showed a prolonged scape latency in LTM at 12wLPS

We evaluated the escape latency in LTM 12wLPS (Fig 5) and observed differences between control *vs.* pristane+LPS (11s *vs.* 140s, *P*=0.016). In this probe, we detected the same behavioral pattern of control mice in relation to the LTM at 7wLPS, in comparison to pristane and pristane+LPS group.

**Fig 5.**
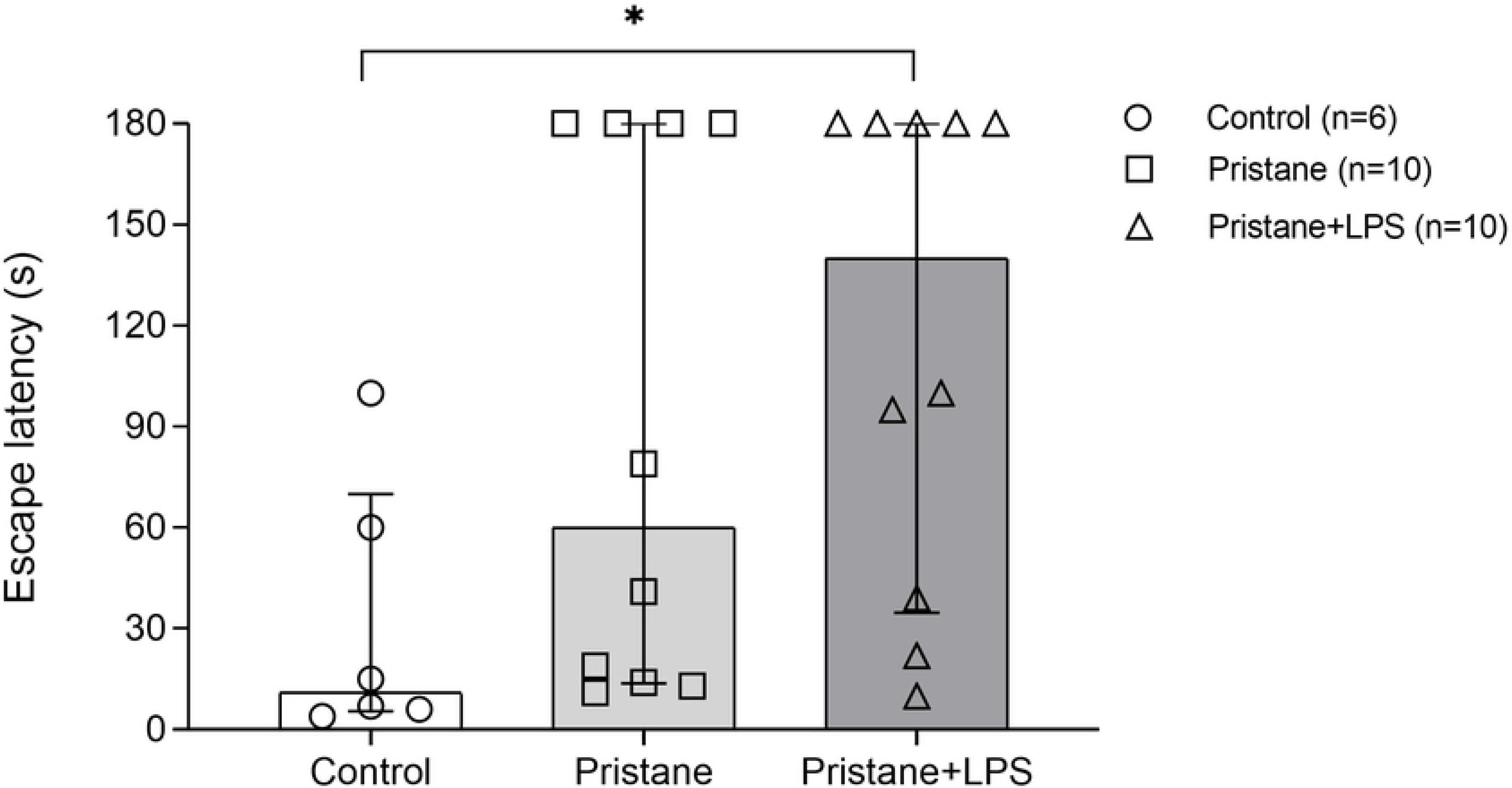
Escape latency of experimental subgroups in LTM at 12wLPS. Total time average in seconds (s) for each mouse to reach and enter to escape box during LTM at 12wLPS. The data were shown in medians and percentile 25 and 75 (P_25_-P_75_). * *P*<0.05.

During LTM at 12wLPS, we did not observe differences between subgroups in the number of errors committed (S4 Fig).

**S4 Fig. Errors in LTM at 12wLPS.** Total errors of subgroups at 12wLPS in LTM. Values are presented as median with percentile 25 and 75 (P_25_-P_75_).

### Pristane and pristane+LPS treated mice produce the highest levels of serum anti-Sm antibodies

We quantified the serum levels of anti-Sm antibodies in experimental groups by ELISA at 7wLPS and 12wLPS. The results were as follows: control group (n=10) 599.9 U/mL (504.9-652.7 U/mL), pristane (n=15) 1617.3 U/mL (1163-2095 U/mL) and pristane+LPS (n=15) 1284.7 U/mL (736.6-2095 U/mL). We observed statistical differences between the control group *vs.* pristane (*P*<0.01), control group *vs.* pristane+LPS (*P*<0.01) but not between pristane *vs.* pristane+LPS (*P*=0.475) (Fig 6, A). To avoid false positives, we calculated the positive index of experimental samples that may be expressed relative to the control values consider as non-immune samples following the manufacture’s recommendations. For this purpose, we calculated the net optical density (OD) + 2 standard deviation (SD) of control samples for obtain the positive index (0.55) and divided each sample net OD by the positive index we obtained differences in the positive index between control *vs.* pristane (0.55 *vs.* 1.21, *P*<0.01) and control *vs.* pristane+LPS (055 *vs.* 1.3, *P*<0.01) (Fig 6, B).

**Fig 6.**
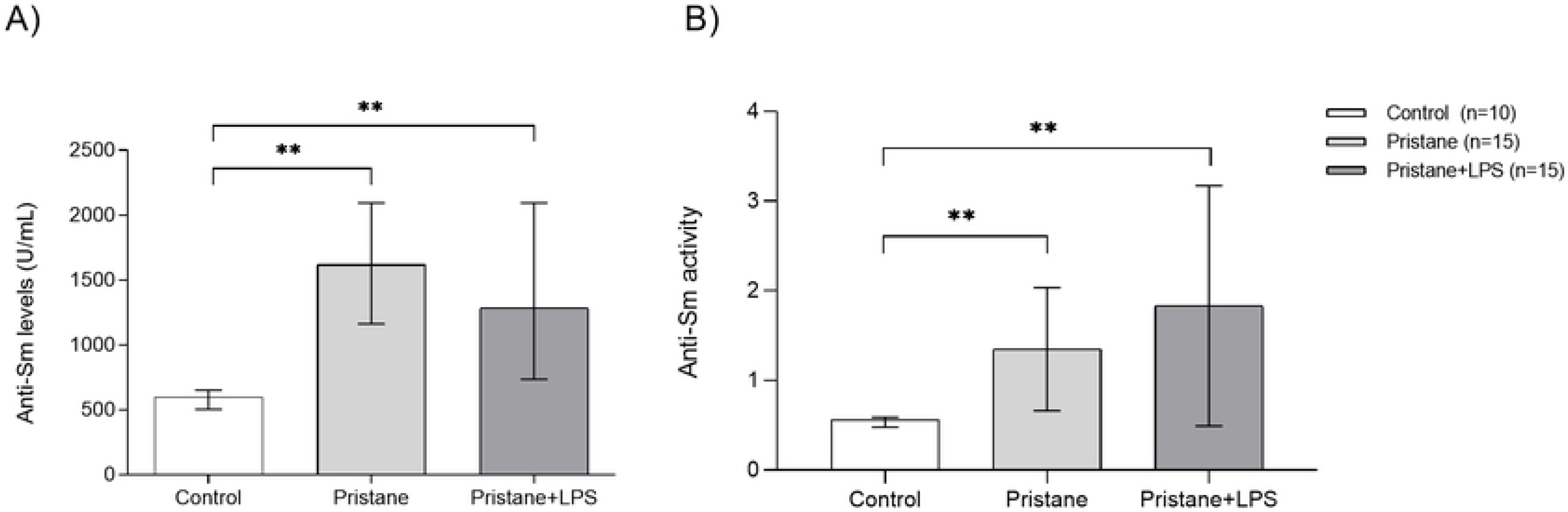
Serum levels of anti-Sm autoantibodies and positive index. A) Serum levels of anti-Sm in experimental subgroups. B) The positive index is shown in arbitrary units. Samples with a value ≥1 were consider positive. \*\**P* <0.01.

### *NR2A* subunit expression decrease in mice treated with pristane and pristane+LPS at 7wLPS and 12wLPS

The expression analysis for hippocampal *NR2A* subunits of the NMDA receptor was evaluated at 7wLPS and 12wLPS (Fig 7). We were able to observe in the pristane group a slight decrease in the relative expression of *NR2A* subunit in 0.51 fold times *vs.* control group (*P*=0.051) and in pristane+LPS a 0.97 fold times *vs.* control group (*P*=0.002). Regarding 12wLPS, we found differences in *NR2A* relative expression between pristane *vs.* control in a 0.95 fold times (*P*=0.004) and in pristane+LPS *vs.* control in a 0.87 fold times (*P*=0.004). It is important to highlight that we did not observe differences in pristane and pristane+LPS groups at 7 wLPS (*P*=0.101) and 12 wLPS (*P*=0.151).

**Fig 7.**
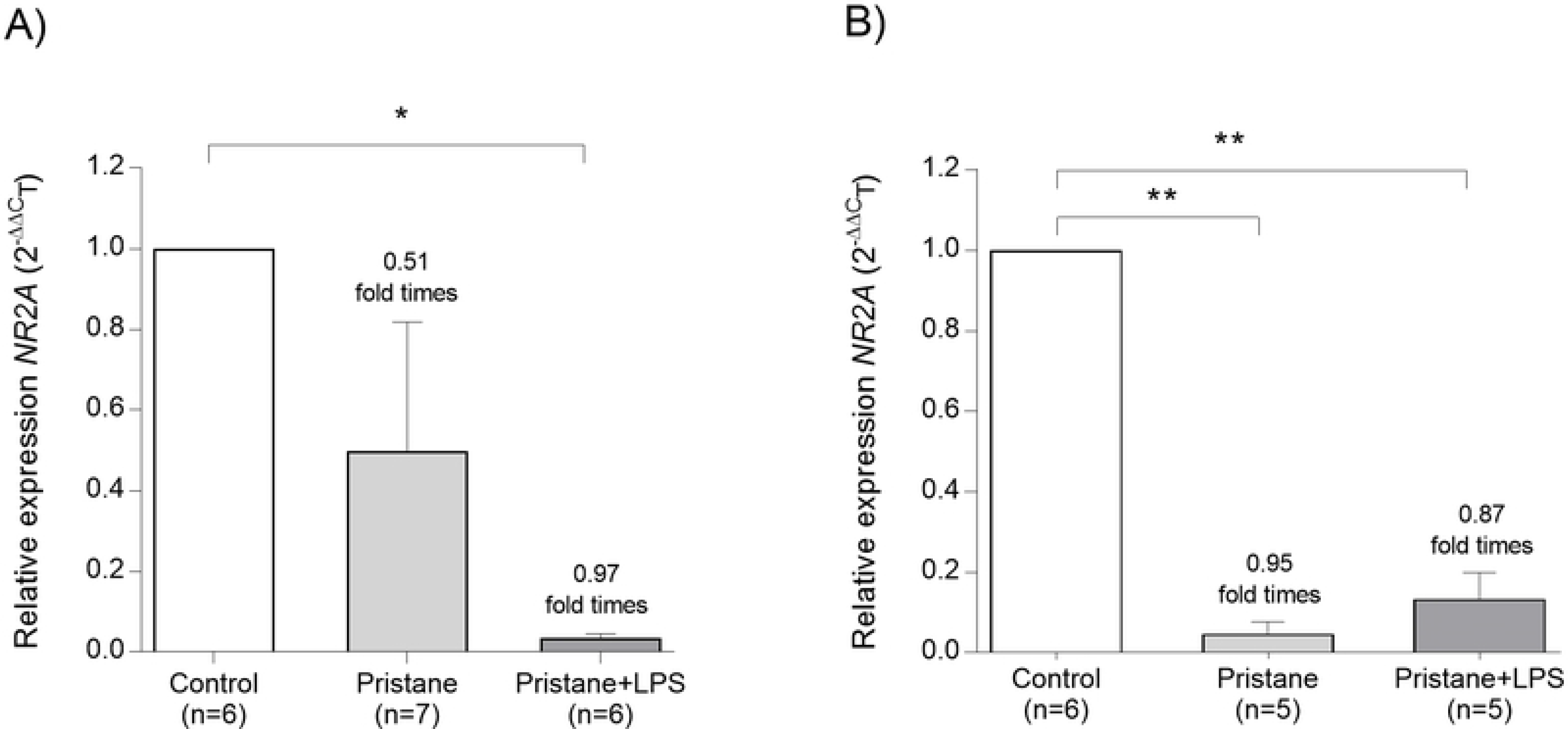
*NR2A* subunit expression of the murine hippocampus at 7wLPS and 12wLPS. Relative expression units were shown in mean (±SEM). * *P*<0.05, ** *P*<0.01.

### *NR2B* subunit expression are not affected than NR2A in mice treated with pristane and pristane+LPS at 7wLPS and 12wLPS

Regarding *NR2B* subunit expression at 7wLPS (Fig 8), we observed in pristane a decrease in 0.68 fold times *vs.* control (*P*=0.001) and in pristane+LPS in 0.41 fold times *vs.* control group (*P*=0.004). In the groups evaluated 12wLPS, we observed in pristane a decrease in 0.56 fold times *vs.* control (*P*=0.004) and in pristane+LPS *vs.* control a 0.36 fold times (*P*=0.004). We did not observe differences between pristane and pristane+LPS groups at 7wLPS (*P*=0.293) and 12wLPS (*P*=0.238).

**Fig 8.**
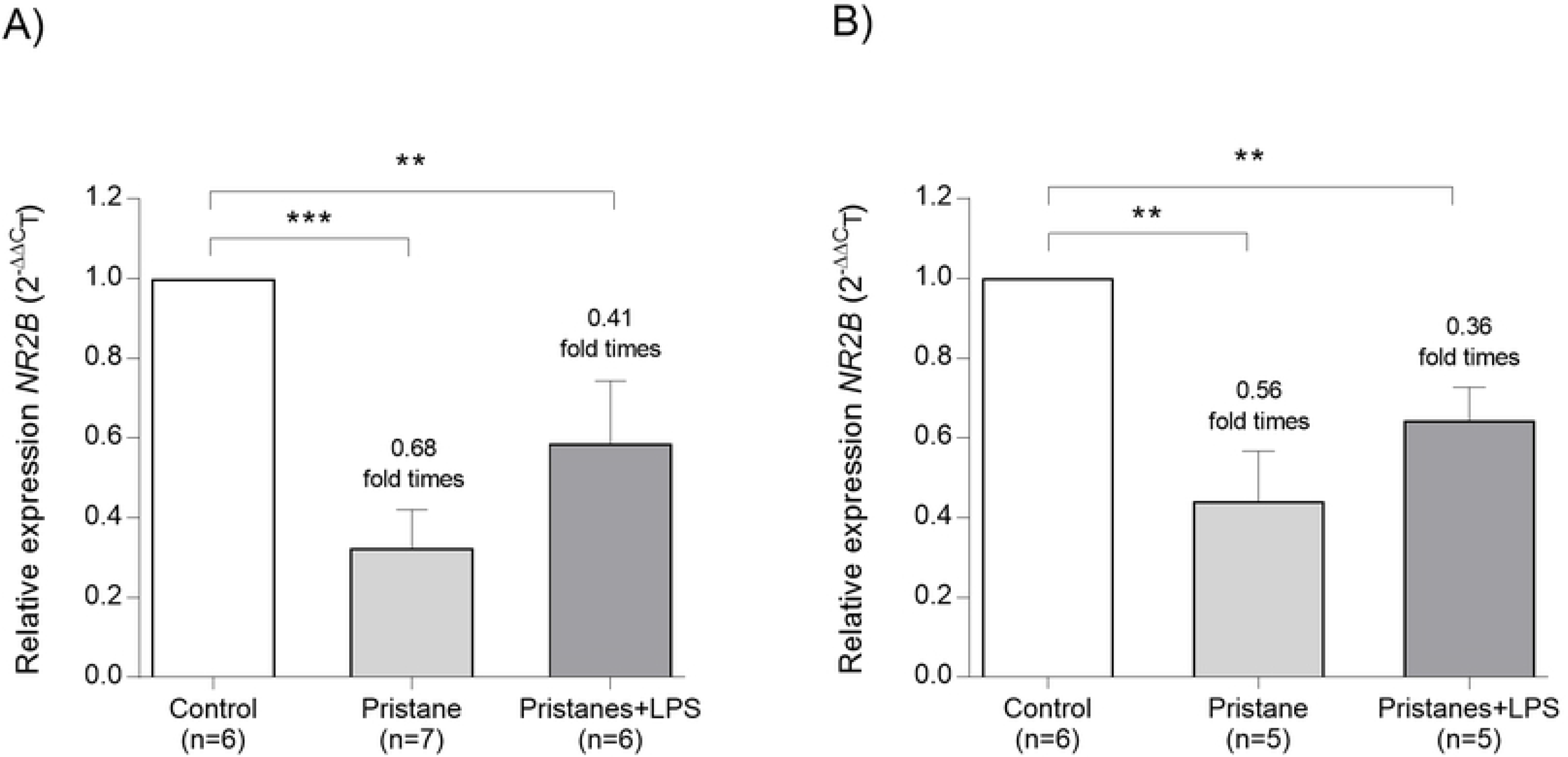
*NR2B* subunit expression of the murine hippocampus at 7wLPS and 12wLPS. Relative expression units were shown in mean (±SEM). \*\**P* <0.01, \*\*\**P*< 0.001.

### *NR2A* subunit expression is affected by anti-Sm antibodies levels in mice treated with pristane and pristane+LPS at 7wLPS and 12wLPS

With the objective of obtain an association between anti-Sm antibodies levels and relative expression of *NR2A/2B* subunits in mice treated with pristane and pristane+LPS at 7wLPS and 12wLPS, we determine a coefficient correlation and obtained an inverse and negative correlation between anti-Sm antibodies and *NR2A* mRNA relative expression (r = −0.461, =0.009; Fig 9, A). Instead, when we analyzed the *NR2B* subunit, we did not observe an association (r = −0.136, *P* = 0.466; Fig 9, B).

**Fig 9.**
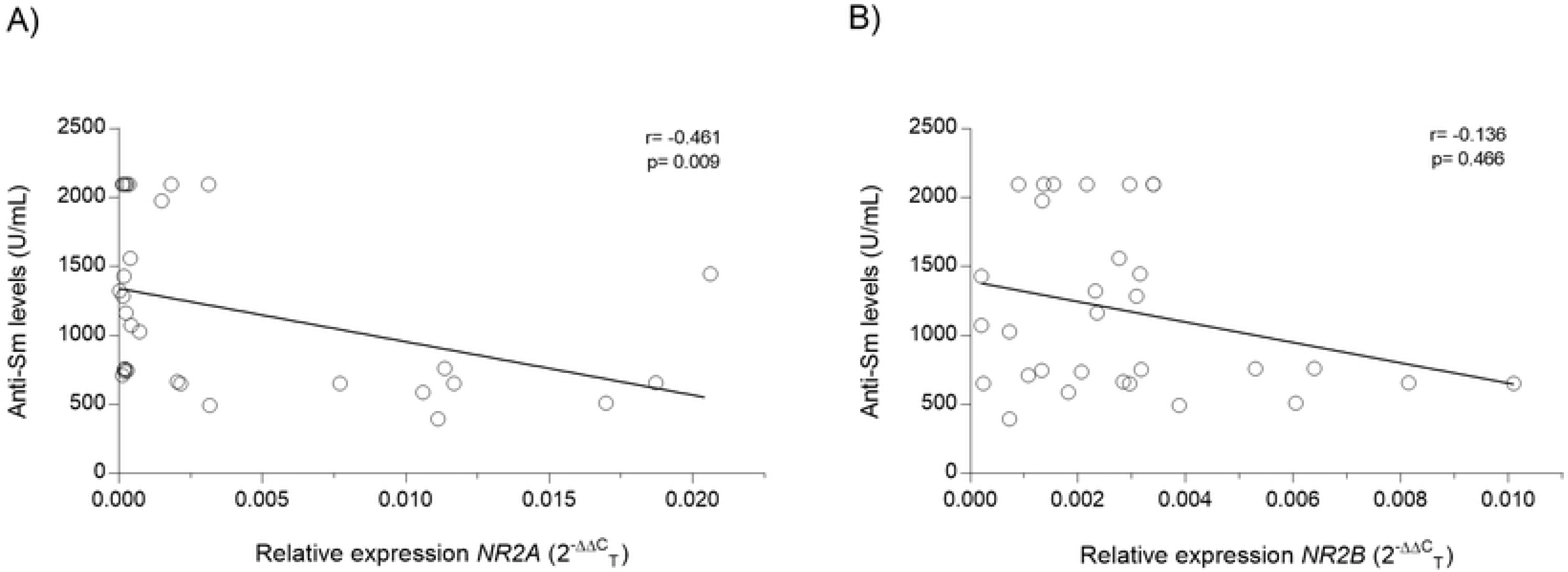
Correlation between anti-Sm antibodies levels and *NR2A/2B* subunit expression. A) Anti-Sm levels/mRNA expression levels of *NR2A* B) Anti-Sm levels/mRNA expression levels of *NR2B*. r= Spearman’s coefficient correlation.

## Discussion

NPSLE is considered a severe condition of SLE physiopathology and the cognitive dysfunction is the more frequent neuropsychiatric alteration with a 15-81% of prevalence [17]. Several studies have proposed that autoantibodies such as anti-dsDNA and anti-Sm produce a cross-reaction with neuronal receptors, attributing a potential pathogenic role in NPSLE [18–20]. One description of this hypothesis was published by Bluestein *et al.* in 1981 where they demonstrated an increased immunoglobulin G (IgG) antineural activity in CSF in SLE patients with active CNS manifestations[21]. These results are in accordance with analysis performed by How *et al.* in 1985, who demonstrated an association between serum antineuronal autoantibodies and NPSLE manifestations[22].

To date, cognitive dysfunction in NPSLE is associated to the presence in serum and CSF of antiphospholipid antibodies and anti-NMDA receptor subunit NR2A/2B (anti-NR2A/2B) antibodies, in addition to disease activity, corticosteroid use, hypertension and chronic damage [3, 17, 23]. Due to the inherent limitations for analyzing the effects of autoantibodies in SLE patients brains, it has been developed murine models of lupus for reproducing and understand the molecular events involved in the induction of excitotoxic neuronal death associated to cognitive impairment in the hippocampal region observed in NPSLE [14]. In this study, we decided to explore the possible cognitive impairment in a murine model that resembles SLE pathogenesis induced by environmental factors with the employ of a hydrocarbon oil pristane. The pristane stimulates in female *BALB/c* mouse the production of proinflammatory cytokines and autoantibodies such as anti-Sm, anti-dsDNA, and anti-U1RNP that in addition with LPS, can disrupt and cross the BBB, altering the permeability of CNS frontier[24]. This phenomenon has been shown in SLE patients through Magnetic Resonance Imaging (MRI) corroborating high levels of permeability of BBB in SLE patients, particularly in hippocampus region more than orbitofrontal, prefrontal, anterior putamen, and globus/thalamus region related to autoantibody production [25].

With the purpose to evaluate the learning and memory process in murine models, behavioral tests are used to assess hippocampal deterioration associated with neuropathologic alterations. The strategic test used to analyze cognitive performance in mice is the Barnes maze, that consists of an elevated circular platform with empty holes and one escape hole around the perimeter. This test takes into advantage the natural preference of rodents for the dark environment and is not influenced by hunger motivation [26]. In this protocol, we used the Barnes maze test in a pristane-lupus induced *BALB/*c mice in two groups: 7wLPS and 12wLPS for STM and LTM divided into experimental subgroups stablished as a control, pristane, pristane+LPS. The Barnes maze protocol considers two basic parameters: the escape latency and errors evaluated in our study during STM (D1-D4) and LTM (D7) at 7wLPS and 12wLPS. In the first probe trial, we observed significant differences in escape latency between pristane/pristane+LPS *vs.* control group at 7wLPS in STM D3-D4, and were maintained in LTM probe. The subgroup of control mice decrease in escape latency as expected, as consequence of memory consolidation and learning process observed in other studies[27]. Nevertheless, mice treated with pristane and pristane+LPS showed an erratic behavior and anxiety, that resulted in a prolongated time to reach and enter in to escape hole. At 12wLPS in STM, we did not observe in pristane and pristane+LPS subgroups the same behavioral pattern that characterize the STM at 7wLPS, in this sense, the mice at 12wLPS showed a decrease in latency to reach the escape hole, evidencing a hippocampal compensatory process [28]. These results show that the pristane have the potential to induce an acute antibody exposure, that can disturb the BBB and, in presence of LPS, these effects can be potentiated in primary stages of the disease, leading for secondary stage the effect of inflammation mediated by T cells and microglial activation with the production of proinflammatory cytokines [29].

Regarding neuronal development and memory consolidation in rodents, studies confirm that the NR2A/B subunits of NMDA mediate certain forms of synaptic plasticity and learning. These receptors are differentially expressed over development with NR2B predominance in mouse brain until NR2A expression increases from the second postnatal week affected by learning and sensory experience[30]. Successful olfactory discrimination learning in rats is associated with an increase in the NR2A/2B ratio and have been proposed that the increase in NR2A stabilize memories [31] and are associated with the emotional behavior regulation in mice [32].

In our study, we quantify the hippocampal mRNA expression levels of *NR2A* and *NR2B* by qRT-PCR and we observed a marked downregulation in the relative expression of *NR2A* more than *NR2B* subunit in 7wLPS and 12wLPS and obtained an inverse and negative correlation between anti-Sm antibodies and *NR2A* mRNA relative expression. This differential effect can be explained by experimental *in vitro* and *in vivo* analysis that relates the presence of autoantibodies (such as anti-NMDA) that affects the *NR2A/2B* synthesis at the nanoscale level and alter the correct receptor function causing synaptic internalization, modifyng the electrical activity. These changes are associated with memory impairment and including the temporary rearrangement of the NR2A/NR2B subunits[33]. On the other hand in relation to anti-Sm antibodies and neuronal receptors, it has been demonstrated that these autoantibodies can disrupt BBB and have a potential neurotoxic effect that is considered prognostic factor for acute confusional state (ACS) in SLE [19, 20], however, more evidence is needed to determine the presence and molecular effect of anti-Sm antibodies on *NR2A/2B* subunits receptors in murine lupus cognitive impairment.

## Conclusions

We conclude that the downregulation of *NR2A/2B* subunits in the pristane-model of lupus in female *BALB/c* mice can be related to anti-Sm autoantibodies production with cognitive impairment consequence in the early stages of autoimmune disease.

## Finantial disclosure

Author(s) received no specific funding for this work.

## Competiting interest

Author(s) have declared that no competing interests exist.

## Author Contributions

### Conceived and experiment designed

Beatriz Teresita Martín-Márquez, Flavio Sandoval-García, Mónica Vázquez-Del Mercado, Yanet Karina Gutiérrez-Mercado and Erika Aurora Martínez-García.

### Performed the experiments

Jonatan Luciano-Jaramillo, Yanet Karina Gutiérrez-Mercado, Rosa Elena Navarro-Hernández, Oscar Pizano-Martínez and Fernanda Isadora Corona-Meraz.

### Analyzed the data

Jonatan Luciano-Jaramillo and Jacinto Bañuelos-Pineda.

### Funding acquisition

Beatriz Teresita Martín-Márquez, Flavio Sandoval-García, Mónica Vázquez-Del Mercado and Jorge Floresvillar-Mosqueda.

### Wrote the paper

Beatriz Teresita Martín-Márquez, Flavio Sandoval-García and Erika Aurora Martínez García.

